# Normalization of single-cell RNA-seq counts by log(x+1)* or log(1+x)*

**DOI:** 10.1101/2020.05.19.100214

**Authors:** A. Sina Booeshaghi, Lior Pachter

## Abstract

Single-cell RNA-seq technologies have been successfully employed over the past decade to generate many high resolution cell atlases. These have proved invaluable in recent efforts aimed at understanding the cell type specificity of host genes involved in SARS-CoV-2 infections. While single-cell atlases are based on well-sampled highly-expressed genes, many of the genes of interest for understanding SARS-CoV-2 can be expressed at very low levels. Common assumptions underlying standard single-cell analyses don’t hold when examining low-expressed genes, with the result that standard workflows can produce misleading results.

**Key Points:**

Lowly expressed genes in single-cell RNA-seq can be easliy misanalyzed.
log(1+x) count normalization introduces errors for lowly expressed genes
The average log(1+x) expression differs considerably from log(x) when x is small
An alternative approach is to use the fraction of cells with non-zero expression

## Results

The *ACE2* receptor, which facilitates entry of SARS-Cov-2 into cells [1], has become one of the most studied genes in the history of genomics over the past two months. There are already hundreds of preprints about the gene (Google Scholar), and it is currently the default gene displayed on the UCSC genome browser [2]. Several studies have reported on the expression of *ACE2* at single-cell resolution, and papers have been rife with speculation about implications of differential *ACE2* mRNA abundance for severity of disease. As is common in single-cell RNA-seq, the expression estimates of *ACE2* are derived from counts that are filtered and normalized. Figure 1a shows an analysis of *ACE2* mRNA in mice lungs (data from [3]). The expression is computed from cells containing at least one copy of the gene. While single-cell RNA-seq expression data has been modeled with many different distributions [4, 5], for simplicity in illustrating our points we model this count data with a simple Poisson random variable *X* with parameter *λ* in order to demonstrate the implications of this restriction. Application of the filter amounts to computing

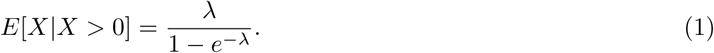

**Figure 1:**
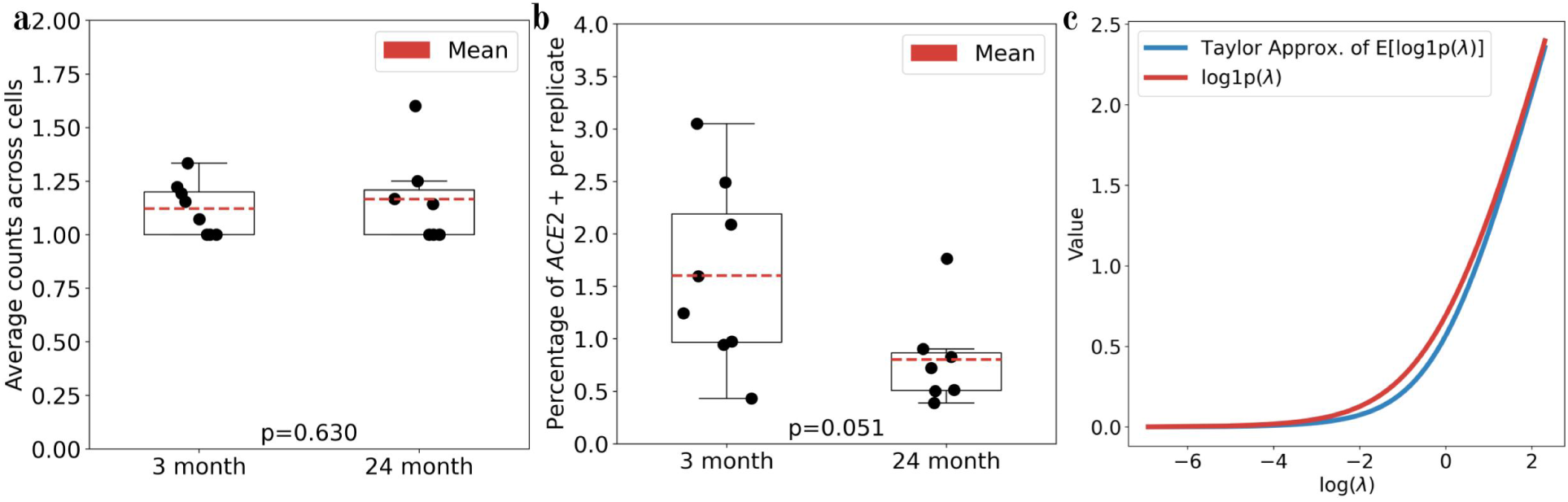
a) Changes in *ACE2* expression in the lungs of eight 3-month old mice and seven 24-month old mice after log1p transformation of the raw counts on the cells with non-zero *ACE2* expression. The p-value was computed using a t-test. b) Changes in *ACE2* expression as determined by the fraction of *ACE2* positive cells. The p-value was computed using a t-test. c) A comparison of the näıve estimate of the expectation of log1p (red) to the Taylor approximation of the expectation of log1p (blue). The code to produce the panels in the figure is available here.

While this is approximately *λ* when *λ* is large, it is close to 1 when *λ* is small [6]. Figure 1b shows the fraction of cells containing at least one copy of *ACE2* [7]. Evidently, Figure 1a creates a misleading impression. While it may appear that average *ACE2* expression is similar between young and old mice, when comparing the fraction of cells with nonzero expression of *ACE2* it is clear that *ACE2* has significantly lower mRNA expression in the lungs of aged mice than young mice.

The fraction *f* of cells with nonzero expression of a gene has a useful statistical interpretation. We leave it as an exercise for the reader to show that the the following estimator for the Poisson rate is consistent:

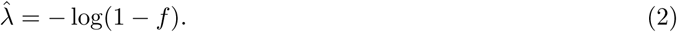

Since *f* is approximately equal to this expression when *f* is small, this provides an interpretation of the fraction of cells with at least one copy of a low-abundance gene as an estimate of the rate parameter *λ* in a Poisson distribution.

Another mistake that we’ve found to be common in reporting *ACE2* expression has to do with the log transformation, frequently used as part of a normalization of counts. Counts are log transformed for two reasons: the first is to stabilize the variance, as the log transform has the property that it stabilizes the variance for random variables whose variance is quadratic in the mean [8, 9]. The rationale of this step for single-cell RNA-seq is manifold: first when performing PCA on the gene expression matrix to find a reduced-dimensional representation that captures the variance, it is desirable that all genes contribute equally. The second rationale for the log transform is that it converts multiplicative relative changes to additive differences. In the context of PCA, this allows for interpreting the projection axes in terms of relative, rather than absolute, abundances of genes.

A seemingly minor technical issue in log transforming counts is that zero counts cannot be “logged”, as log(0) is undefined. To circumvent this problem, it is customary to add a “pseudocount”, e.g. +1, to each gene count prior to log transforming the data [10]. We denote this by log1p (see units of Figure 1a), in accordance with nomenclature standard in scientific computing [11]. For a gene with an average of *λ* counts where *λ* is large, it is intuitive that the average of the log1p transformed counts is approximately log(*λ*). However, this is not true for small *λ*. An understanding of the result of applying the log1p transform begins with the observation that for a random variable *X*, *E*[*f* (*X*)] is not, in general, equal to *f* (*E*[*X*]). For example, if *X* is a Poisson random variable with parameter *λ*, it is not true that *E*[log(1 + *X*)] = log(*λ* + 1). By Taylor approximation,

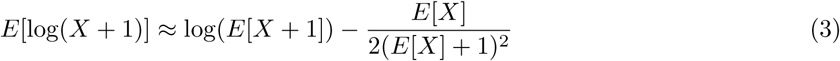

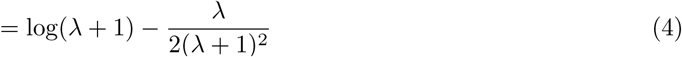

This shows that for low-expressed genes, the average log1p expression differs considerably from log(*λ*) (see Figure 1c). Thus, while a 2-fold change for large *λ* translates to a log(2) difference after log1p, that is not the case for small *λ*.

In summary, while single-cell RNA-seq atlases offer detailed information about the transcriptomic profiles of distinct cell types, their use to examine specific genes, as has been done recently in the study of SARS-CoV- 2 infection related genes, requires care. Methods should not be used unless their limitations are understood. For example, while it doesn’t matter whether one uses log(x+1) or log(1+x), the filtering and normalization applied to counts can affect comparative estimates in non-intuitive ways. Moreover, there are subtle problems that arise when working with small counts that transcend the elementary issues we have raised [12, 13]. These matters are not theoretical; we leave the identification of published preprints and papers that have ignored the issues we’ve raised, and hence reported misleading results, as another exercise for the reader.

## Acknowledgments

We thank Charles Herring, Michael Hoffman, Harold Pimentel, Jeffrey Spence, and Valentine Svensson for helpful comments.

